# A GluN2B disease-associated variant promotes degradation of NMDA receptors via autophagy

**DOI:** 10.1101/2025.01.12.632651

**Authors:** Taylor M. Benske, Marnie P. Williams, Pei-Pei Zhang, Adrian J. Palumbo, Ting-Wei Mu

**Author notes:** Corresponding author: Ting-Wei Mu.

## Abstract

N-methyl-D-aspartate receptors (NMDARs) are essential for excitatory neurotransmission and their pathogenic variants can lead to proteostasis defects and thus neurological diseases. However, how the proteostasis network degrades pathogenic variants is not well understood. Here, we demonstrated that the R519Q GluN2B variant is retained in the endoplasmic reticulum (ER) and fails to traffic to the surface to form functional NMDARs. Pharmacological and genetic inhibition of autophagy results in the accumulation of this variant, indicating that it is degraded by the autophagy-lysosomal proteolysis pathway. Since GluN2B has a cytosolic LIR motif, which can interact with cytosolic autophagy machinery, we demonstrated that disrupting this LIR motif impairs the autophagic clearance of this variant. Additionally, the R519Q variant is recognized by ER-phagy receptors, including CCPG1 and RTN3L. Our result provides the molecular mechanism for the degradation of NMDAR variants and identifies a pathway for targeted therapeutic intervention for neurological disorders with dysfunctional NMDARs.

**Summary:** NMDA receptors are essential for excitatory neurotransmission and their proteostasis defects lead to neurological diseases. Benske et al. report that pathogenic R519Q variants predispose GluN2B subunits to degradation and clearance by the autophagy-lysosomal pathway.

## Introduction

*N*-methyl-D-aspartate receptors (NMDARs) are ionotropic glutamate receptors that mediate excitatory neurotransmission, synaptic development and plasticity, and maintain the excitation-inhibition balance in the central nervous system (Hansen et al., 2021). These heterotetrameric receptors assemble from two obligatory, glycine-binding, GluN1 subunits and two GluN2(A-D) or GluN3(A-B) subunits, encoded by seven *GRIN* genes (Traynelis et al., 2010). In the human forebrain, NMDARs predominantly contain GluN2A and/or GluN2B subunits, likely comprising triheteromeric receptors (GluN1/GluN2A/GluN2B) (Chazot and Stephenson, 1997; Hansen et al., 2014; Rauner and Köhr, 2011; Tovar et al., 2013). Structurally, GluN subunits consist of four distinct domains: an extracellular amino-terminal domain (ATD), the ligand-binding domain (LBD) which is formed by two segments S1 and S2, the transmembrane domain (TMD), consisting of three helices and one reentrant loop, and an intracellular carboxyl-terminal domain (CTD) (**Figure 1A**) (Karakas and Furukawa, 2014; Paoletti et al., 2013). The glutamate-binding GluN2 subunits impart unique biophysical and pharmacological properties to the NMDARs contributing to their distinct regional and temporal expression patterns (Sanz-Clemente et al., 2013; Vieira et al., 2020), where the GluN2B subunit is most abundantly expressed prenatally/embryonically, while GluN2A is predominately expressed at postnatal, mature synapses (Akazawa et al., 1994).

**Figure 1.**
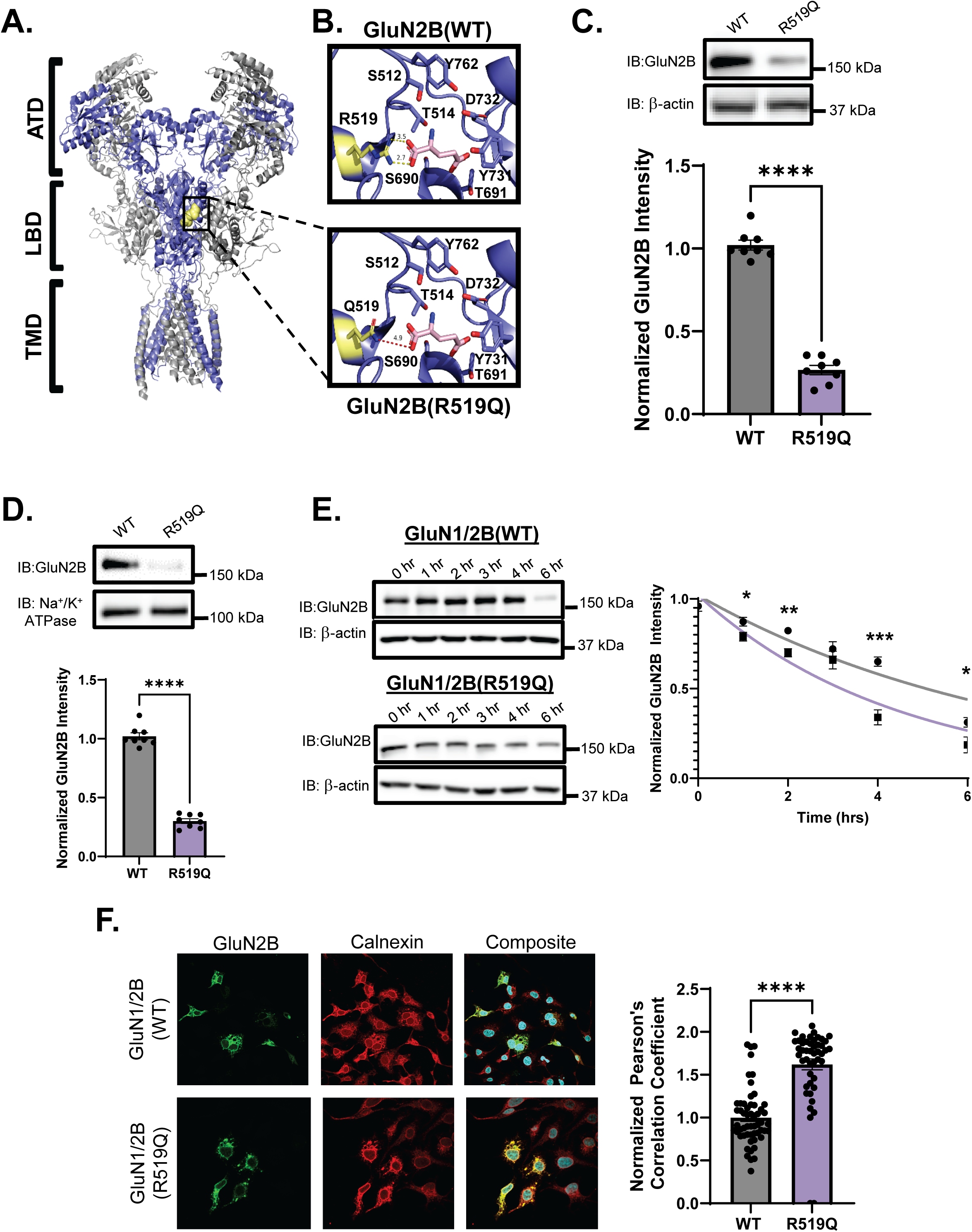
Molecular characterization of the disease-associated variant GluN2B_R519Q. **(A)** Structure of the rat tetrameric GluN1_GluN2B NMDA receptor in complex with glutamate, with the GluN1 subunit in gray, and the GluN2B subunit in blue (PDB:9ARI) (Chou et al., 2024). The amino-terminal domain (ATD), ligand-binding domain (LBD), and transmembrane domain (TMD) are shown in ribbon format with the R519 residue selected for study in yellow spheres. **(B)** Residues involved in the glutamate binding site of the WT GluN2B subunit are represented a sticks, with the glutamate ligand in pink (upper). Mutagenesis of the R519 residue to a glutamine eliminates the electrostatic interaction with glutamate (lower). **(C)** Effects of R519Q on total GluN2B protein expression levels 48 hrs post transient transfection of HEK293T with GluN1 and GluN2B constructs at a 1:1 ratio to express WT or GluN2B_R519Q variant NMDARs (n=7). β-actin serves as the soluble total protein loading control. (**D)** Surface biotinylation assay to monitor the influence of the R519Q DAV on the surface expression of NMDARs 48 hrs post transient transfection. Na^+^/K^+^ ATPase served as a membrane protein loading control (n=8). **(E)** HEK293T cells stably expressing WT or R519Q GluN2B NMDARs were subjected to cyclohexmide (100 µg/mL) for the indicated times to determine the stability and rate of degradation (n=5). **(F)** Immunofluorescence images of GluN2B (green) and calnexin (red) an ER marker to assess ER accumulation (scale bar = 20 μm). Pearson’s coefficients are reported (n>30) and statistical significance was determined using a Mann-Whitney test. All data normalized to the appropriate loading control and data are presented as mean ± SEM. Statistical significance was determined using an unpaired two-tailed Student’s t-test between two groups or an analysis of variance (ANOVA) followed by a post-hoc Dunnett’s test for comparison in multiple groups. Significance level defined as *p<0.05, **p<0.01, ***p<0.001, ****p<0.0001.

Whole genome sequencing has identified numerous disease-associated-variants (DAVs) within the *GRIN* genes in patients with neurodevelopmental disorders including autism spectrum disorder, intellectual disabilities, developmental delays, and epilepsies (Benke et al., 2021; Burnashev and Szepetowski, 2015; Platzer et al., 2017; XiangWei et al., 2018). Patients often present additional symptoms including hypotonia, movement disorders, behavioral abnormalities, and gastrointestinal issues (ClinVar). Despite this, there are currently minimal treatment options for GRIN disorders, and many that do specifically target NMDAR dysfunction require the receptors to be expressed on the cell surface (Benke et al., 2021). *GRIN* DAVs are typically *de novo* and heterozygous, in which loss-of-function mutations exhibit haploinsufficiency (Benke et al., 2021; XiangWei et al., 2018). DAVs within the GluN2B subunit are clustered in the LBD and the ion pore, underscoring their critical role in receptor activity (Amin et al., 2021). Residual variation intolerance scores obtained through analyzing healthy populations for genetic variation, found that *GRIN* genes, exhibit a lower than anticipated frequency of single-nucleotide polymorphisms, indicating that mutations are likely to lead to disease states (Benske et al., 2022). Remarkably, *GRIN2B* was found to be amongst the top 1.07% of intolerant genes within the human genome and have a high probability of being loss-of-function intolerant score (Karczewski et al., 2020; Petrovski et al., 2013). Underscoring the importance of the GluN2B subunit, mouse models with homozygous knockdown of the GluN2B subunit or homozygous truncations of the CTD results in neonatal and perinatal lethality respectively, that cannot be rescued by substitution with GluN2A subunits (Hamada et al., 2014; Kutsuwada et al., 1996; Sprengel et al., 1998). Despite this, the *GRIN* genes are highly polymorphic, with over 4000 reported mutations identified, in which 445 variants have been characterized to be pathogenic. 95% of GRIN variants are found within the GluN1, GluN2A, and GluN2B subunits (Santos-Gómez et al., 2022).

Numerous studies have been devoted to exploring the electrophysiological consequences of DAVs in NMDARs, particularly in heterologous expression systems (Li et al., 2019; Platzer et al., 2017; Swanger et al., 2016; Vyklicky et al., 2018) and more recently animal models have been developed to investigate the biological impact of select DAVs (Candelas Serra et al., 2024; Shin et al., 2020; Sullivan et al., 2024). Despite this, a majority of the DAVs remain uncharacterized. Altered surface trafficking and membrane localization due to NMDAR DAVs can contribute to the pathophysiology in disease states, as proper localization at postsynaptic sites is essential for receptor function, though the precise mechanism by which DAVs influence surface expression of NMDARs remains poorly understood. NMDAR subunits are folded and assembled within the endoplasmic reticulum (ER) to form functional heterotetramers and must pass stringent quality control checks prior to their anterograde transport to the Golgi apparatus (Hanus and Ehlers, 2016; Horak et al., 2014; Jeyifous et al., 2009; Lichnerova et al., 2015; Setou et al., 2000). The ER is an intracellular organelle that utilizes chaperone proteins to assist in the folding, assembly, glycosylation, and formation of disulfide bonds of nascent polypeptides in order to promote their native confirmation (Adams et al., 2019). Multiple studies have demonstrated that ligand binding is an essential quality control check within the ER that is essential for the forward trafficking of NMDARs, indicating that the structural integrity of the LBD is vital (Kenny et al., 2009; Netolicky et al., 2024; She et al., 2012; Swanger et al., 2016). The presence of missense mutations can exacerbate the inefficiency of functional folding and assembly (Gershenson et al., 2014), which can lead to the accumulation of misfolded proteins within the ER, resulting in ER stress and activation of the unfolded protein response (UPR) (Balchin et al., 2016; Wu and Kaufman, 2006; Yang et al., 2024). Recently, work from our group reported the capacity of BIX, a potent BiP activator, to adapt the proteostasis network of NMDARs containing GluN2A DAVs and restore their deficient folding, trafficking, and function, while also preventing their degradation (Zhang et al., 2024). Despite this, the specific details governing the quality control and protein homeostasis of NMDARs in regards to their folding, assembly, trafficking, and degradation remain poorly understood (Benske et al., 2022).

Misfolded, aggregated, and short-lived proteins are degraded by ER-associated degradation (ERAD), in which the protein substrates are retrotranslocated into the cytosol and degraded via the ubiquitin proteasome system. ER-phagy (reticulophagy) is an alternative degradation pathway in which large bulk proteins, aggregates, and ERAD-resistant populations are degraded via the selective autophagy of the ER sheets, tubules, and localized cargo, by the autophagy lysosomal pathway (Sun and Brodsky, 2019; Wu and Rapoport, 2018). Membrane bound ER-phagy receptors sense imbalances within the ER lumen and link the ER to the autophagy machinery via the interaction of ATG8 family proteins, LC3b/GABARAP and their LC3b interacting regions (LIR) motifs. These autophagy cargo receptors include FAM134 paralogs (A,B,C), RTN3L, ATL3, SEC62, CCPG1, and TEX264 (Dikic, 2018; Wilkinson, 2020). These receptors are located to distinct regions of the ER, where FAM134b is restricted to the curvature of ER sheets, RTN3L, ATL3 and TEX264 are within the ER tubules, and CCPG1 and SEC62 are concentrated in the ER sheets after ER stress near insoluble proteins. Ultimately, the ER-phagy receptors mediate cargo segregation, membrane shaping, and clearance of specific regions of the ER through the lysosomal degradation pathway.

While we have extensive knowledge regarding the general effects of mutations within the GluN2B polypeptide sequences, less is understood about how the resulting aberrant proteins engage with the degradation machinery that assist in their catabolic clearance. Therefore, in this study we sought to understand how NMDAR subunits utilize the proteolytic pathways for their clearance and degradation. Here, we used NMDARs containing GluN2B subunits with the R519Q variant in the LBD to investigate the pathogenic influence on their proteostasis networks using HEK293T cells exogenously expressing NMDARs. Our results indicated that autophagy plays a critical role in regulating the clearance of pathogenic NMDARs.

## Results

### The R519Q GluN2B variant reduces the protein expression and stability of NMDARs and results in ER retention

Considering that a majority of DAVs are located within the LBD, which plays a crucial role in ER quality control of NMDARs, this study focuses on the proteolytic degradation of the R519Q variant within the LBD of the GluN2B subunit (**Figure 1A, 1B**) (Chou et al., 2024). The R519Q variant is reported in ClinVar as likely pathogenic and associated with intellectual disability, and yet, to the best of our knowledge, it has not been extensively characterized in the literature. To evaluate the effects of the R519Q variant on the protein levels of NMDARs, we co-expressed wild type (WT) or R519Q GluN2B subunits with GluN1 subunits in HEK293T cells, using a 1:1 ratio of cDNA. Cell cultures were supplemented with MK-801, a potent NMDAR antagonist, to prevent glutamate-induced excitotoxicity, and harvested 48 hours after transfection. Western blot analysis of total proteins revealed that the R519Q expression was reduced drastically, with a 5.0-fold decrease compared to WT GluN2B (**Figure 1C**). Given that the surface expression of NMDARs is critical for their physiological function, we carried out a surface biotinylation assay and revealed a 5.0-fold decrease in the GluN2B surface expression of the R519Q variant (**Figure 1D**).

The R519 residue has been shown to directly interact with the glutamate ligand, forming a salt bridge that is critical for maintaining agonist potency and stability of the protein-ligand complex (**Figure 1B**, upper) (Chou et al., 2024). The R519Q missense mutation likely destroys the salt bridge (**Figure 1B**, lower), destabilizing the complex and causing protein misfolding and excessive degradation. Next, we performed a cycloheximide (CHX) chase assay in order to determine the half-lives of the WT and R519Q variant GluN2B subunit and assess their stability. HEK293T cells stably expressing NMDARs containing either WT or R519Q GluN2B subunits were treated with CHX, a potent inhibitor of protein synthesis, and chased for the indicated time. Quantification via an exponential decay fit showed that the half-life of the R519Q variants was significantly reduced, 3.439 hr, compared to the WT, 5.898 hr, GluN2B subunit (**Figure 1E**). This reduced stability of the R519Q variant correlates with their steady-state protein levels observed earlier and indicates faster degradation kinetics.

NMDAR subunits must properly fold and assemble within the ER before they can traffic to the plasma membrane. Therefore, since the R519Q variant demonstrated a profound decrease in surface expression of the GluN2B subunit, we asked whether this variant was retained in the ER. We used immunocytochemistry confocal imaging to determine the colocalization of the WT and R519Q variant with the endogenous ER-resident protein calnexin. The R519Q variant displayed substantial overlap and exhibited a greater Pearson’s correlation coefficient compared to the WT GluN2B subunit (**Figure 1F**). This indicates that the R519Q variant was retained within the ER, likely failing quality control measures and preventing their incorporation into function receptors expressed on the plasma membrane. Collectively, our results indicate that the R519Q variant perturbs NMDAR proteostasis by reducing the stability of the GluN2B subunit resulting in ER retention and reduced total and surface expression, which may underlie the pathophysiology of *GRIN2B*-related neurodevelopmental disorders.

### The R519Q DAV drives GluN2B degradation towards autophagy and clearance via the ER-phagy receptors CCPG1 and RTN3

Because we demonstrated the R519Q variant has reduced stability and increased ER retention, we sought to elucidate how the proteolytic pathways effectively facilitate its clearance. We pharmacologically inhibited lysosomal degradation with bafilomycin A1 (Baf-A1), a potent V-type ATPase inhibitor, and proteasomal degradation with MG132, a potent proteasome inhibitor, in HEK293T cells stably expressing NMDARs with WT or R519Q GluN2B. In WT-expressing cells, treatment with both Baf-A1 and MG132 resulted in a modest accumulation of GluN2B, 1.75-fold, compared to the DMSO control, suggesting that both pathways are utilized (**Figure 2A**, left). In contrast, cells expressing the R519Q NMDARs displayed a striking 4.2-fold increase upon treatment with Baf-A1, while treatment with MG132 did not significantly affect GluN2B expression (**Figure 2A**, right). These findings represent a 2.4-fold higher accumulation of the R519Q variant subunit relative to the WT after Baf-A1 treatment, suggesting that the R519Q variant predisposes the GluN2B subunit to the clearance via autophagy (**Supplemental Figure 1A**). In order to confirm that the observed accumulation of the R519Q variant following Baf-A1 treatment was a direct result of autophagic inhibition, we used additional pharmacological modulators of autophagy to monitor the effects on the GluN2B subunit (**Figure 2B**). HEK293T cells stably expressing NMDARs containing R519Q variants were treated for 24 hrs with autophagy inhibitors and activators. Inhibition of autophagy with 3-methyladednine (3-MA; 50 mM), a PI3K inhibitor for VPS34, resulted in a 3.1-fold increase in total GluN2B levels, similar to the Baf-A1 treatment (20 nM) (**Figure 2C**). Conversely, autophagy was activated with rapamycin (100 nM), a specific inhibitor of mTOR, or by SMER28 (10 μM), a small molecule that induces autophagy in an mTOR-independent fashion, and neither significantly altered the GluN2B expression (**Figure 2C**). Interestingly, surface biotinylation assays revealed that 24 hr treatment with 3-MA increased surface expression of R519Q variants, while Baf-A1, rapamycin, and SMER28 did not influence the surface expression (**Supplemental Figure 1B**).

**Figure 2:**
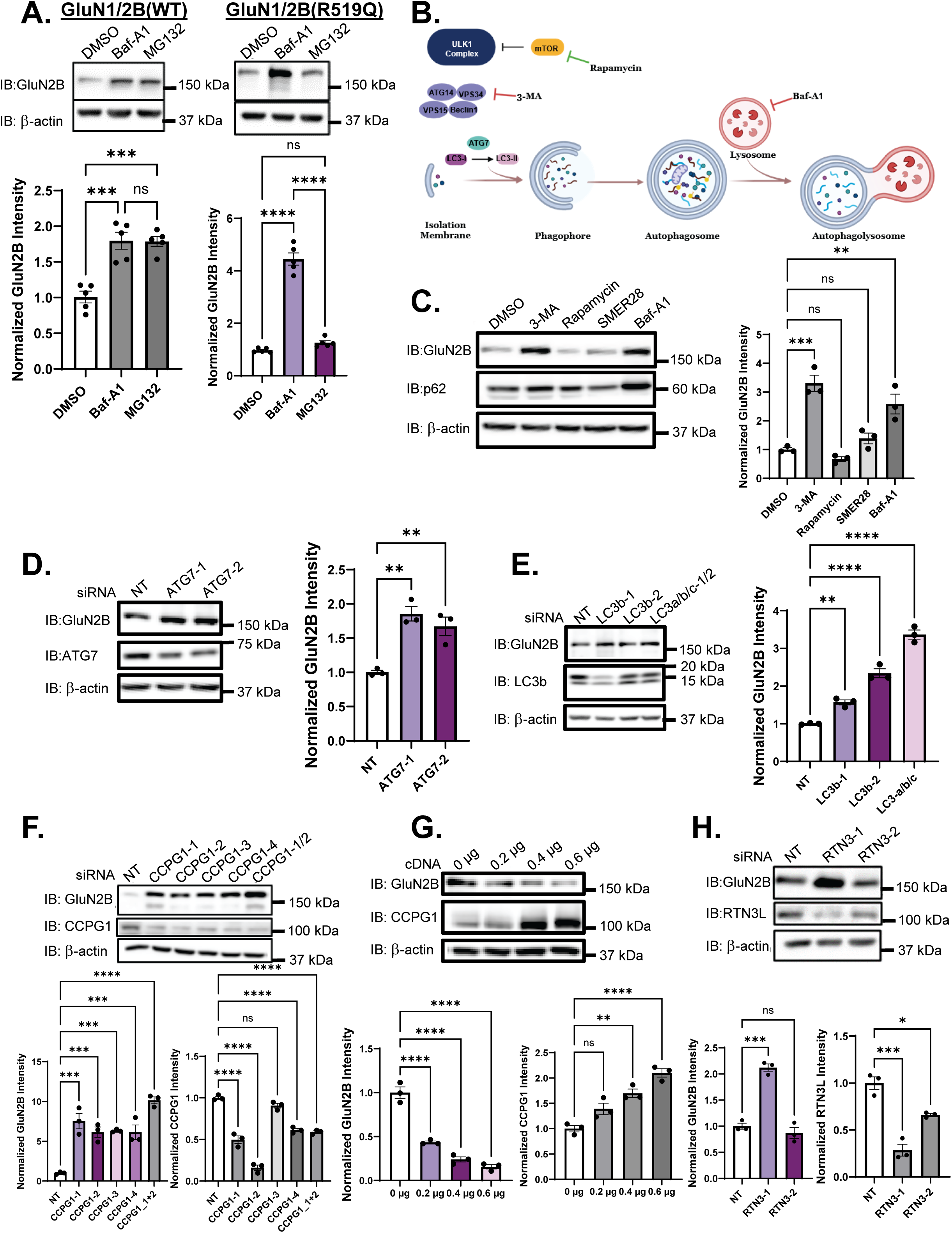
The GluN2B subunit undergoes degradation via autophagy. **(A)** Inhibition of the proteasome with MG132 (10 µM) and the lysosome with Bafilomycin-A1 (1 µM) for 6 hr effect on the GluN2B subunit in HEK293T cells stably expressing recombinant WT or R519Q NMDARs (n=5). β-actin serves as the soluble total protein loading control. **(B)** Schematic of macroautophagy pathway, including induction, nucleation and formation of the isolation membrane, elongation and autophagosome formation, and fusion with the lysosome forming the autophagolysosome. Rapamycin is shown inhibiting mTOR which results in autophagy activation, while 3-MA and Baf-A1 inhibit autophagy at different stages as shown. **(C)** HEK293T cells stably expressing R519Q NMDARs were treated with autophagy activators (SMER28 10 µM, Rapamycin 100 nM) and inhibitors of autophagy (3-MA 50 mM and Baf-A1 20 nM) for 24 hours. Changes to GluN2B expression were monitored via immunoblot and p62 was used as a marker for autophagic flux (n=3). **(D)** siRNA knockdown of ATG7 effect on R519Q variant GluN2B expression and immunoblot was performed 48hrs after knockdown (n=3). Non-targeting siRNA was used as a control for each condition, and two independent siRNA constructs were used to knockdown gene expression. Knockdown efficiency was determined by looking at protein levels of the knockdown target. β-actin served as the soluble total protein loading control. **(E)** siRNA knockdown of LC3b effects on R519Q variant GluN2B expression. Immunoblot performed 48hrs after knockdown (n=3). Non-targeting siRNA was used as a control for each condition, and two independent siRNA constructs were used to knockdown gene expression. Knockdown efficiency was determined by looking at protein levels of the knockdown target. **(F)** siRNA transfection on HEK293T cells stably expressing R519Q NMDARs with siRNA targeting CCPG1. Immunoblot performed 48hrs after knockdown (n=4). Non-targeting siRNA was used as a control and four independent siRNA constructs were used to knockdown gene expression. β-actin served as the soluble total protein loading control. **(G)** Overexpression of CCPG1 cDNA in HEK293T cells stably expressing GluN2B_R519Q NMDARs. cDNA dose response of CCPG1 was performed. Cells were harvested 24hrs after transient transfection (n=3). β-actin served as the soluble total protein loading control. **(H)** RTN3L siRNA transfection of HEK293T cells stably expressing R519Q NMDARs Immunoblot performed 48hrs after knockdown, probed for RTN3L and RTN3s (n=3). Non-targeting siRNA was used as a control for each condition, and two independent siRNA constructs were used to knockdown gene expression. Knockdown efficiency was determined by looking at protein levels of the knockdown target. β-actin served as the soluble total protein loading control. All data normalized to the appropriate loading control and data are presented as mean ± SEM. Statistical significance was determined using unpaired two-tailed Student’s t-test between two groups or an analysis of variance (ANOVA) followed by a post-hoc Tukey test for comparison in multiple groups. Significance level defined as p<0.05; *p<0.05, **p<0.01, ***p<0.001, ****p<0.0001.

Additionally, we determined the effect of inhibiting autophagy machinery genetically on the clearance of the R519Q variant. ATG7 is an essential E1-like enzyme required for the formation of the autophagosome by conjugating ATG8 (LC3b and GABARAP) family proteins to phosphatidylethanolamine to form the isolation membrane (**Figure 2B**). ATG7 knockdown with two unique siRNAs resulted in increased GluN2B levels in cells stably expressing the R519Q variant, further supporting autophagy in its degradation (**Figure 2D**, **Supplemental Figure 1C**). Consistently, knocking down LC3b increased expression of the GluN2B subunit (**Figure 2E**, **Supplemental Figure 1D**).

Furthermore, given that the R519Q variant is retained in the ER and preferentially targeted to the lysosome for degradation, we determined the potential role of ER-phagy in their degradation. Remarkably, knocking down CCPG1 using siRNAs, a selective ER-phagy receptor within the ER sheets, induced a substantial increase in R519Q variant expression (**Figure 2F**). Consistently, overexpressing CCPG1 led to a dose-dependent decrease in the R519Q variant levels in HEK293T cells (**Figure 2G**), confirming that CCPG1 mediates the autophagic clearance of the R519Q variants from the ER. Additionally, knockdown of RTN3L, another ER-phagy receptor, also caused a significant 2.0-fold increase in the R519Q variant expression (**Figure 2H**). Together, these results establish that the R519Q GluN2B subunit is primarily degraded through lysosomal pathways, with CCPG1 and RTN3L playing critical roles in facilitating its clearance from the ER.

### Genetic inhibition of autophagy limits GluN2B degradation and results in the accumulation of the R519Q DAV

ATG7 is a critical autophagy component that initiates the formation and elongation of the autophagosome through the lipidation of ATG8 homologs (**Figure 2B**). To further examine the role of autophagy in the degradation of NMDARs, we utilized HEK293T cells with a CRISPR-mediated knockout (KO) of ATG7 to overexpress NMDARs containing WT or R519Q variant GluN2B subunits. The ATG7 KO allowed us to study the sustained effects of chronic autophagy depletion on GluN2B proteostasis and account for any autophagy degradation driven by transient transfection-induced cell stress. We first validated the ATG7 KO line using immunoblot assay to confirm the near-complete depletion of ATG7 protein (**Figure 3A**). As expected, there were significantly elevated protein levels of p62, a canonical autophagy substrate, indicating that autophagic flux was impaired, confirming the functionality of this model for our investigations. To assess the impact of autophagy deficiency on GluN2B stability, we exogenously expressed NMDARs containing WT or R519Q variant GluN2B subunits in the ATG7 KO HEK293T cells. Remarkably, immunoblot analysis revealed that the R519Q variant GluN2B subunit was expressed at levels comparable to the WT in the absence of autophagy (**Figure 3B**). We next sought to determine whether the surface expression of the R519Q variant could be rescued in ATG7 KO HEK293T cells. Intriguingly, surface biotinylation assays demonstrated that the R519Q variant was only able to traffic to the plasma membrane around 20% of the WT (**Figure 3C**). These findings indicate that inhibiting autophagy is not sufficient to allow the accumulated R519Q variant pool in the ER to assemble into functional receptors for their anterograde trafficking.

**Figure 3:**
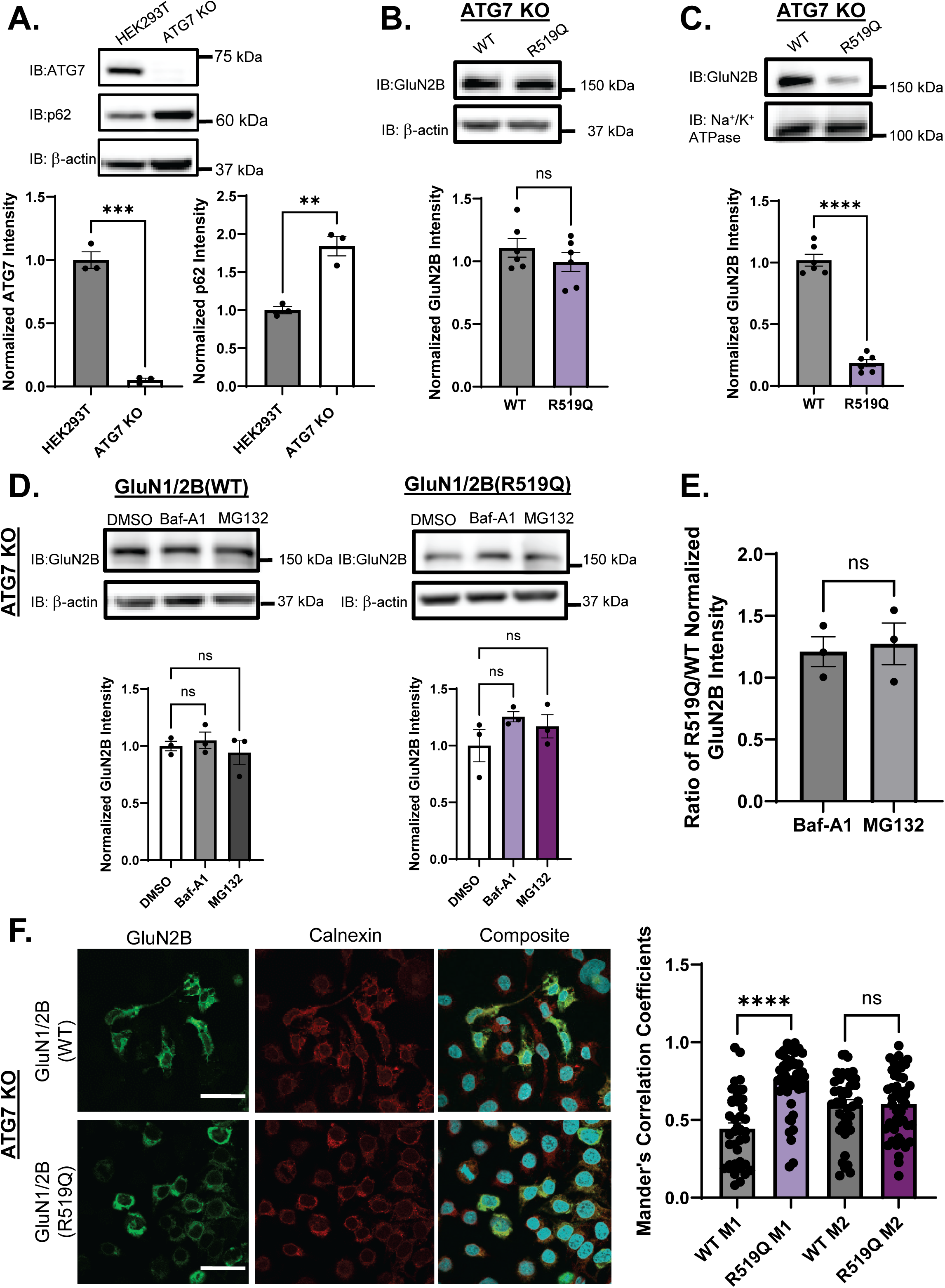
Genetic depletion of Autophagy in ATG7 KO lines rescues R519Q total expression. **(A)** Validation of HEK293T ATG7 KO line. Immunoblot was used to quantify levels of ATG7 in HEK293T and ATG7 KO lines. p62 was used to assess impaired autophagic flux (n=3). **(B)** Effects of R519Q on total GluN2B protein expression levels 48 hrs post transient transfection with GluN1 and GluN2B constructs at a 1:1 ratio in ATG7 KO HEK293T to express WT or GluN2B_R519Q variant NMDARs (n=6). β-actin served as the soluble total protein loading control. **(C)** Surface biotinylation assay to monitor the influence of the R519Q DAV on the surface expression of NMDARs 48 hrs post transient transfection (n=6). Na^+^/K^+^ ATPase served as a membrane protein loading control. **(D)** Inhibition of the proteasome with MG132 (10 µM) and the lysosome with Baf-A1 (1 µM) for 6 hrs on the effect of the GluN2B subunit in ATG7 KO HEK293T cells stably expressing recombinant WT or R519Q NMDARs (n=3). β-actin served as the soluble total protein loading control. **(E)** Ratio of the normalized accumulation of R519Q/WT GluN2B upon treatment with Baf-A1 and MG132. **(F)** Immunofluorescence images of GluN2B (green) and calnexin an ER marker (red) to assess ER accumulation. Mander’s coefficients are reported, where M1= ratio of GluN2B/Calnexin and M2= ratio of Calnexin/GluN2B (scale bar: 20 μm, n>=30) and statistical significance was determined using a Mann-Whitney test. All data normalized to the appropriate loading control and data are presented as mean ± SEM. Statistical significance was determined using an unpaired two-tailed Student’s t-test between two groups or an analysis of variance (ANOVA) followed by a post-hoc Tukey test for comparison in multiple groups. Significance level defined as *p<0.05, **p<0.01, ***p<0.001, ****p<0.0001.

Because the ATG7 KO cells lack macroautophagy but not all lysosomal or proteasomal degradation, we next investigated whether there was compensation in the degradation of the GluN2B subunit. Thereby, we expressed WT and R519Q variant NMDARs in the ATG7 KO lines. 48 hrs after transient transfection we inhibited the lysosome by treatment with Baf-A1 (1 μM, 6 hr) and the proteasome by treatment with MG132 (10 μM, 6 hr). SDS-PAGE and immunoblot analysis showed no significant accumulation of either the WT or the R519Q variant upon inhibition of either proteolytic pathway (**Figure 3D, 3E**). These results suggest that there is no compensation by the proteasome to facilitate the degradation of GluN2B subunits, nor is there a significant fraction of NMDARs degraded by lysosome via endocytosis or chaperone-mediated autophagy.

Given that we were unable to see changes to surface expression, we determined where the GluN2B receptor subunits were accumulating within the cell. Immunofluorescence confocal microscopy was used to examine the localization of the GluN2B subunits with the ER marker calnexin. Colocalization analysis using Mander’s coefficients M1 revealed an increase in the GluN2B/calnexin colocalization of the R519Q variant in comparison to the WT (**Figure 3F**). The Mander’s coefficients M2 of the calnexin/GluN2B colocalization were comparable between the WT and the R519Q variant. Taken together, these findings demonstrate that autophagy plays a critical role in the degradation of GluN2B, particularly in the presence of the R519Q variant. Furthermore, while inhibition of autophagy increases total GluN2B levels, it does not promote the surface expression of the R519Q variant, underscoring the importance of understanding the pathogenic defects of DAVs on the biogenesis and the effects on the trafficking of NMDAR DAVs.

### The highly conserved GluN2B LC3b-interacting region (LIR) motif influences the proteostasis of NMDA receptors

LC3b-interacting region (LIR) motifs are present in proteins that undergo selective autophagy and have been observed in autophagy cargo receptors, basal autophagy proteins, proteins associated with vesicles, and signaling proteins and receptors that are degraded by selective autophagy. LIR motifs are characterized by the consensus sequence [W/F/Y]X_1_X_2_[L/I/V] (**Figure 4A**), where the first residue is a bulky aromatic residue, X_1_ and X_2_ can be any amino acid and the fourth residue is a hydrophobic residue (Birgisdottir Å et al., 2013; Rogov et al., 2023). We utilized the iLIR database, an online web resource for identifying LIR motif-containing proteins in eukaryotes (Jacomin et al., 2016). We discovered that the human GluN2B subunit (Uniprot Accession No: Q13224) was predicted to contain an F-type LIR motif within the CTD consisting of the residues DTFVDL at positions 1305-1310 (**Figure 4A, 4B**). Notably, in F-type LIR domains the N or C terminus must be flanked by an acidic residue, further the residue present at X_1_ must be Val, Cys, Ile, or Glu. These constraints are in place as F-type LIR motifs demonstrate weaker affinity for the ATG8 proteins and the higher amount of electrostatic interactions has been shown to make up for lower affinities (Birgisdottir Å et al., 2013; Wirth et al., 2019). Of all the GluN subunits, the GluN2B subunit alone has a predicted canonical LIR motif. Although the CTD of the GluN2A and GluN2B subunit are elongated allowing for similar protein interactions and modulations, such as phosphorylation via CAMKII and interaction with PSD95 (Bard et al., 2010; Barria and Malinow, 2005; Doré et al., 2014), a sequence alignment of the human GluN2B and GluN2A CTDs in the region containing the LIR motif showed that while 50% of this region is conserved, the analogous sequence of the GluN2A (DNIVDK at position 1293-1298) does not fit the characteristics of an LIR motif (**Figure 4B**). Additionally, the GluN2B LIR motif is absolutely conserved across mammalian species including humans, mice, rats, dogs, and chimpanzees, emphasizing its functional importance to the GluN2B subunit (**Figure 4C**).

**Figure 4:**
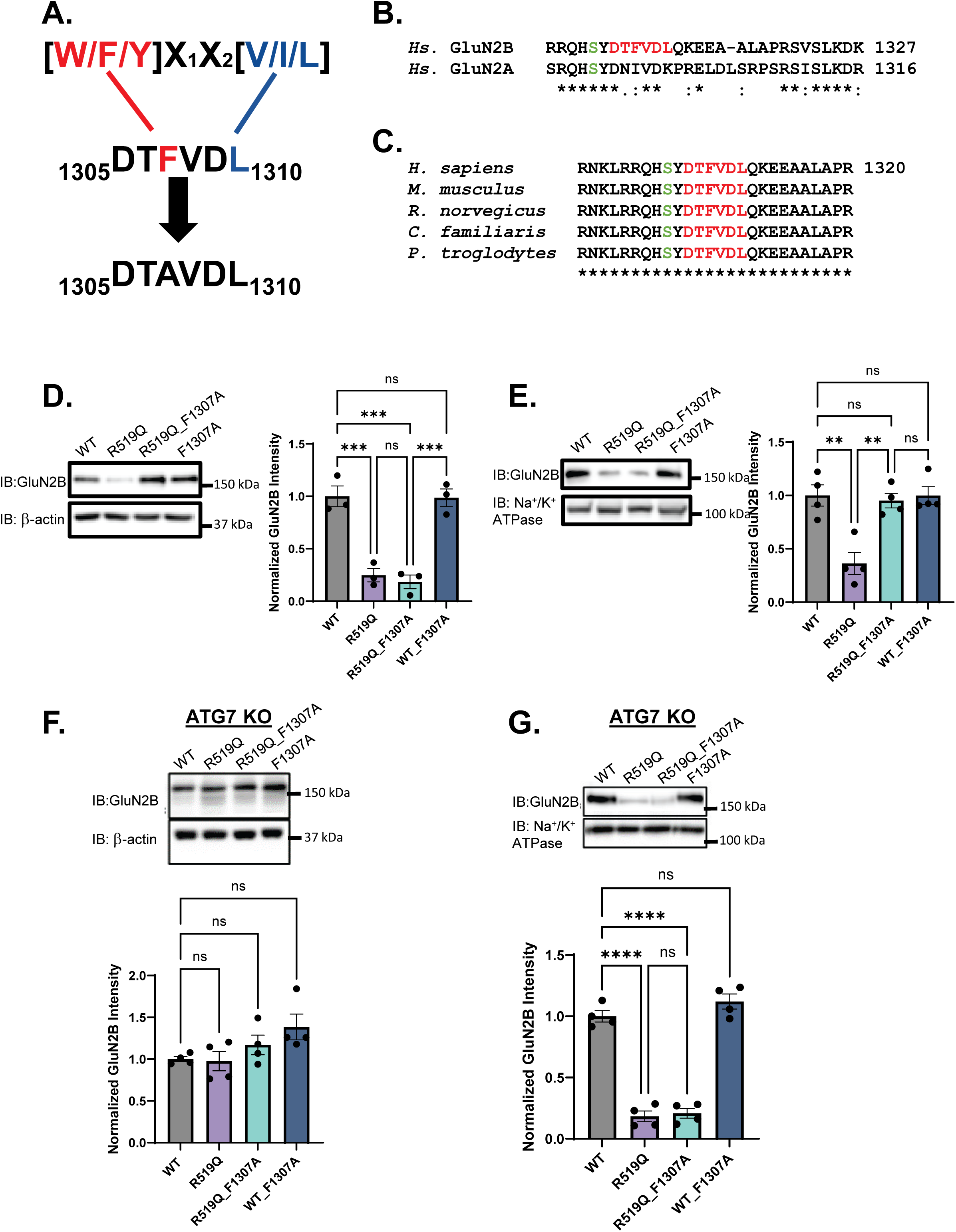
The GluN2B LC3-interacting region motif is highly conserved. (**A**) Canonical LC3b interacting motif consensus sequence pointing to the LIR motif identified in the GluN2B subunit and subsequent alanine substitution. Aromatic residue shown in red, hydrophobic residue in blue. **(B)** Sequence alignment of *Homo sapiens* GluN2A and GluN2B CTD amino acid sequence in the region of interest. The LIR motif is in red in the GluN2B subunit. The serine residue that is phosphorylated by CAMKII is shown in green for each subunit. **(C)** Amino acid sequence alignment of mammalian species demonstrating species conservation of the GluN2B CTD region of interest surrounding the LIR motif. **(D)** Effects of LIR motif disruption on WT, WT_F1307A and R519Q, R519Q_F1307A total GluN2B protein expression levels 48 hrs post transient transfection of HEK293T cells transfected with a 1:1 ratio of GluN1 and GluN2B constructs to express WT or GluN2B_R519Q variant NMDARs (n=4). β-actin served as the soluble total protein loading control. **(E)** Surface biotinylation assay to monitor the influence of the LIR domain disruption on the WT and R519Q DAV surface expression of NMDARs 48 hrs post transient transfection (n=3). Na^+^/K^+^ ATPase served as a membrane protein loading control. **(F)** Effects of LIR motif disruption on total expression of GluN2B subunits expressed in ATG7 KO cells. 48 hrs post transient transfection of GluN1 and GluN2B constructs with a 1:1 ratio of GluN1 and GluN2B constructs to express WT, WT_F1307A, R519Q, R519Q_F1307A variant NMDARs (n=4). β-actin served as the soluble total protein loading control. **(G)** Surface biotinylation assay to monitor the influence of the LIR motif disruption on WT and the R519Q variant GluN2B subunits on the surface expression of NMDARs in ATG7 KO cells 48 hrs post transient transfection (n=4). Na^+^/K^+^ ATPase served as a membrane protein loading control. All data normalized to the appropriate loading control and data are presented as mean ± SEM. Statistical significance was determined using an unpaired two-tailed Student’s t-test between two groups or an analysis of variance (ANOVA) followed by a post-hoc Tukey test for comparison in multiple groups. Significance level defined as *p<0.05, **p<0.01, ***p<0.001, ****p<0.0001.

To investigate the role of the LIR motif, we performed alanine substitution at the critical phenylalanine (F1307) residue (**Figure 4A**) to disrupt the hydrophobic interaction required for binding to ATG8 proteins. We co-expressed GluN1 with either WT, WT_F1037A, R519Q, or R519Q_F1307A GluN2B subunits in HEK293T cells. Western blot analysis demonstrated that the R519Q_F1307A disruption of the LIR domain increased the total expression of the GluN2B subunit to levels comparable to the WT (**Figure 4D**). However, the WT_F1307A disruption of the LIR motif did not influence the total expression of the GluN2B subunit when compared to the WT. These results indicate that the LIR domain positively regulates the autophagic clearance of aberrant GluN2B subunits and prevents their accumulation. We then assessed the surface expression via surface biotinylation assays to determine whether the accumulation of the R519Q subunits by LIR motif disruption was able to restore their surface expression. As expected, the surface expression of the R519Q_F1307A variant displayed equivalent surface expression to the R519Q variant alone (**Figure 4E**), indicating that the trafficking defects of the R519Q variant cannot be rescued by disrupting the LIR domain alone. In addition, the WT_F1307A did not reduce the surface level compared to WT (**Figure 4E**).

To further evaluate the role of the LIR motif in the autophagy-dependent degradation of the GluN2B subunits, we expressed the WT and R519Q LIR variants in ATG7 KO HEK293T cells. Immunoblot analysis revealed similar expression of all GluN2B subunits (**Figure 4F**), further supporting that autophagy is essential in the removal of the R519Q variant. However, when we tested whether the disruption of the LIR motif would influence the surface expression in the ATG7 KO line, we discovered that the R519Q_F1307A variant was expressed on the surface at equal levels to the R519Q variant (**Figure 4G**). These findings indicate that the LIR motif in the GluN2B subunit is crucial for the autophagic clearances of misfolded proteins, but does not significantly alter receptor surface trafficking.

### Disruption of the LIR motif attenuates the lysosomal degradation of the R519Q DAV

To examine whether disruption of the LIR motif influences the sensitivity of the GluN2B subunit to degradation, we inhibited the lysosome and proteasome using Baf-A1 (1 μM, 6 hr) and MG132 (10 μM, 6 hr) respectively in HEK293T cells co-expressing GluN1 with either WT, WT_F1037A, R519Q, or R519Q_F1307A GluN2B subunits. In the case of WT GluN2B and WT_F1307A LIR disruption, there was equal accumulation with inhibition of either the lysosome or the proteasome with a 1.7-fold accumulation in the WT upon inhibition of either pathway and a similar increase in the WT_F1307A (**Figure 5A**). Assessment of the ratios of GluN2B in response to proteolytic inhibition shows no significant change between the WT_F1307A LIR disruption and the WT GluN2B subunits (**Figure 5C** right). In contrast, the R519Q_F1307A variant exhibited a 1.6-fold increase in response to both lysosomal and proteasomal degradation inhibition. This contrasts the R519Q variant that exhibits a 4.1-fold accumulation upon inhibition of the lysosome but no accumulation upon inhibition of the proteasome (**Figure 5B**). Quantitative analysis determined that the R519Q_F1307A exhibited a 60% reduction in accumulation in response to Baf-A1 treatment compared to the R519Q variant. Conversely, the R519Q_F1307A variant displays 160% accumulation of the GluN2B upon treatment with MG132 when compared to the R519Q variant (**Figure 5D** right). These results demonstrate a shift towards proteasomal degradation when the LIR domain of the R519Q variant is disrupted, which resembles the degradation patterns observed in the WT_F1307A.

**Figure 5:**
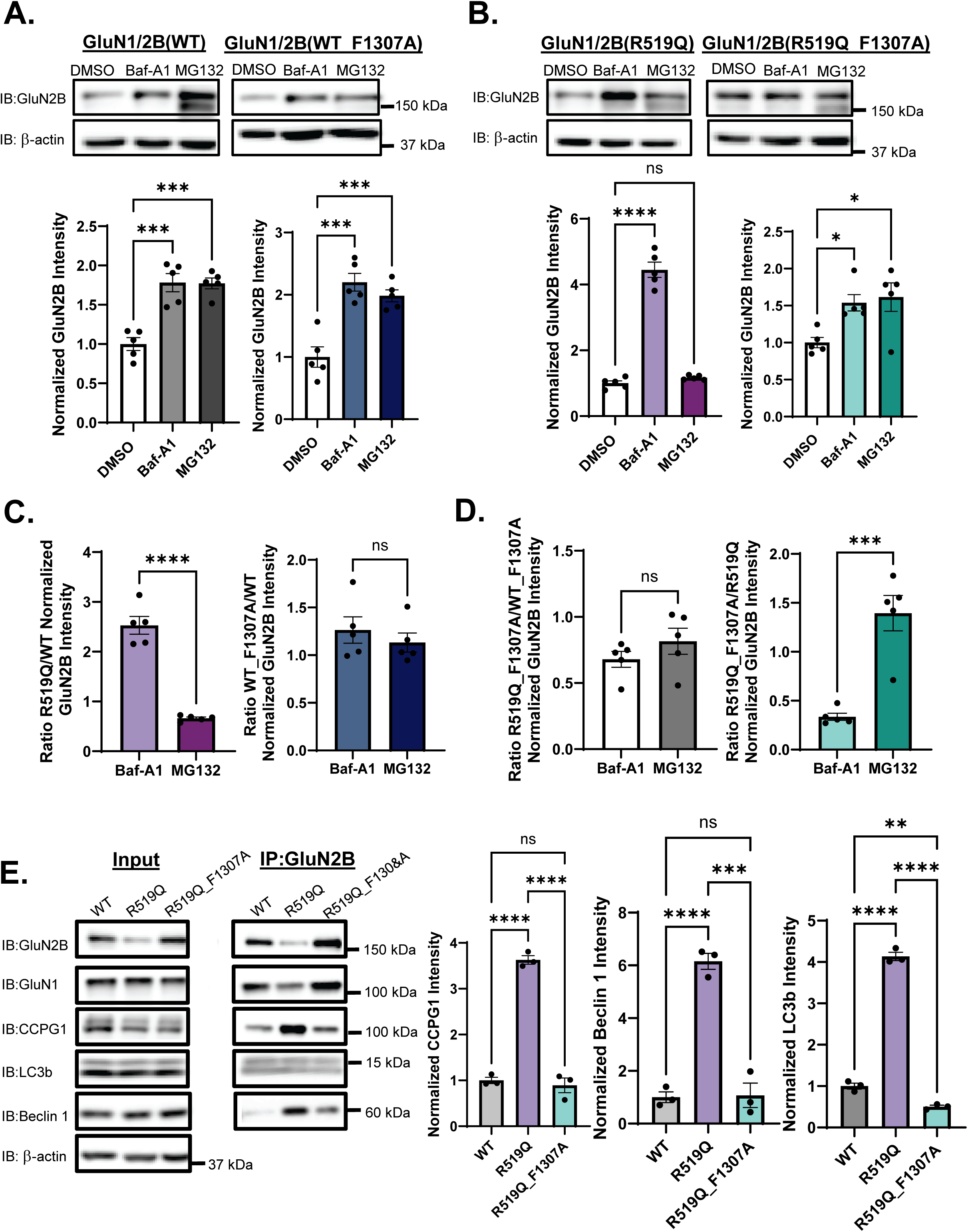
Effects of LIR motif disruption on GluN2B autophagic degradation. **(A)** Inhibition of the proteasome with MG132 (10 µM) and the lysosome with Baf-A1 (1 µM) for 6 hrs effect on the WT (left) and WT_F1307A (right) GluN2B subunit in HEK293T cells 48hrs after transient transfection (n=5). β-actin served as the soluble total protein loading control. **(B)** Inhibition of the proteasome with MG132 (10 µM) and the lysosome with Baf-A1 (1 µM) for 6 hrs effect on the R519Q (left) and the R519Q_F1307A (right) GluN2B subunit in HEK293T cells 48hrs after transient transfection (n=5). β-actin served as the soluble total protein loading control. **(C)** Ratio of the normalized accumulation of R519Q/WT GluN2B (left) and the ratio of the normalized accumulation of WT_F1307A/WT GluN2B (right) upon treatment with Baf-A1 and MG132. **(D)** Ratio of the normalized accumulation of R519Q_F1307A/WT_F1307A GluN2B (left) and the ratio of the normalized accumulation of R519Q_F1307A/R519Q GluN2B (right) upon treatment with Baf-A1 and MG132. **(E)** Co-immunoprecipitation of HEK293T cells exogenously expressing WT, R519Q, or R519Q_F1307A NMDARs with GluN2B pull down and autophagy proteins and ER-phagy receptors. Cells were treated with Baf-A1 (20 nM for 24 hrs), 24 hrs post transient transfection to enrich autophagy proteins for interaction (n=3). All data normalized to the appropriate loading control and data are presented as mean ± SEM. Statistical significance was determined using an unpaired two-tailed Student’s t-test between two groups or an analysis of variance (ANOVA) followed by a post-hoc Tukey test for comparison in multiple groups. Significance level defined as *p<0.05, **p<0.01, ***p<0.001, ****p<0.0001.

Finally, we examined whether disruption of the LIR motif compromises interactions between GluN2B and proteins critical for autophagic degradation. Co-immunoprecipitation assays were performed using a GluN2B antibody to pull-down HEK293T cell lysates overexpressing WT, R519Q, or R519Q_F1307A GluN2B subunits. Cells were treated with Baf-A1 (1 μM, 24 hr) in order to enrich autophagy proteins and enhance detection of potential interactions. Probing for key autophagy proteins including CCPG1, Beclin 1, and LC3b revealed notable differences in their interaction profiles (**Figure 5E**). Most notably, while the R519Q_F1307A was able to interact with CCPG1 and Beclin 1, the interaction was substantially decreased compared to the R519Q variant. These results suggest that the presence of the R519Q variant promotes the interaction with autophagy proteins when compared to the WT subunit, and that the LIR motif facilitates the key interactions implicated in the autophagic degradation of the R519Q variant.

## Discussion

Here our collective data characterizes the effect of the pathogenic R519Q variant present in the LBD of the GluN2B subunit on the proteostasis of NMDARs. Our results support the mechanism of selective degradation of variant GluN2B subunits from the ER as proposed in **Figure 6**. GluN2B subunits containing DAVs within the LBD display reduced expression and trafficking to the cell surface. This is the result of the retention of misfolded proteins within the ER, as they fail quality control measures that ensure ligand binding. As a result, these mutations are targeted towards degradation by selective autophagy through interaction with the ER-phagy receptors in the ER sheets via CCPG1 and within ER tubules via RTN3L. CCPG1 contains 3 identified cargo interacting regions (CIR) that are present within the ER lumen that have demonstrated the ability to identify misfolded and accumulated cargo (Ishii et al., 2023). In the case of CCPG1 and RTN3L, they contain LIR motifs that have been demonstrated to recruit autophagic machinery and initiate the formation of the autophagosome membrane. Furthermore, the LIR motif within the CTD of the GluN2B influences the degradation of aberrant subunits and disruption diminishes the interaction with autophagy proteins, including with the ER-phagy receptors. However, our study did not elucidate whether this was through direct interactions or loss of interactions through a mediator or adaptor protein. It is unclear what proteins engage with NMDAR subunits within the ER to mediate their biogenesis and target them for degradation, warranting further investigations into the proteostasis networks of NMDAR subunits.

**Figure 6:**
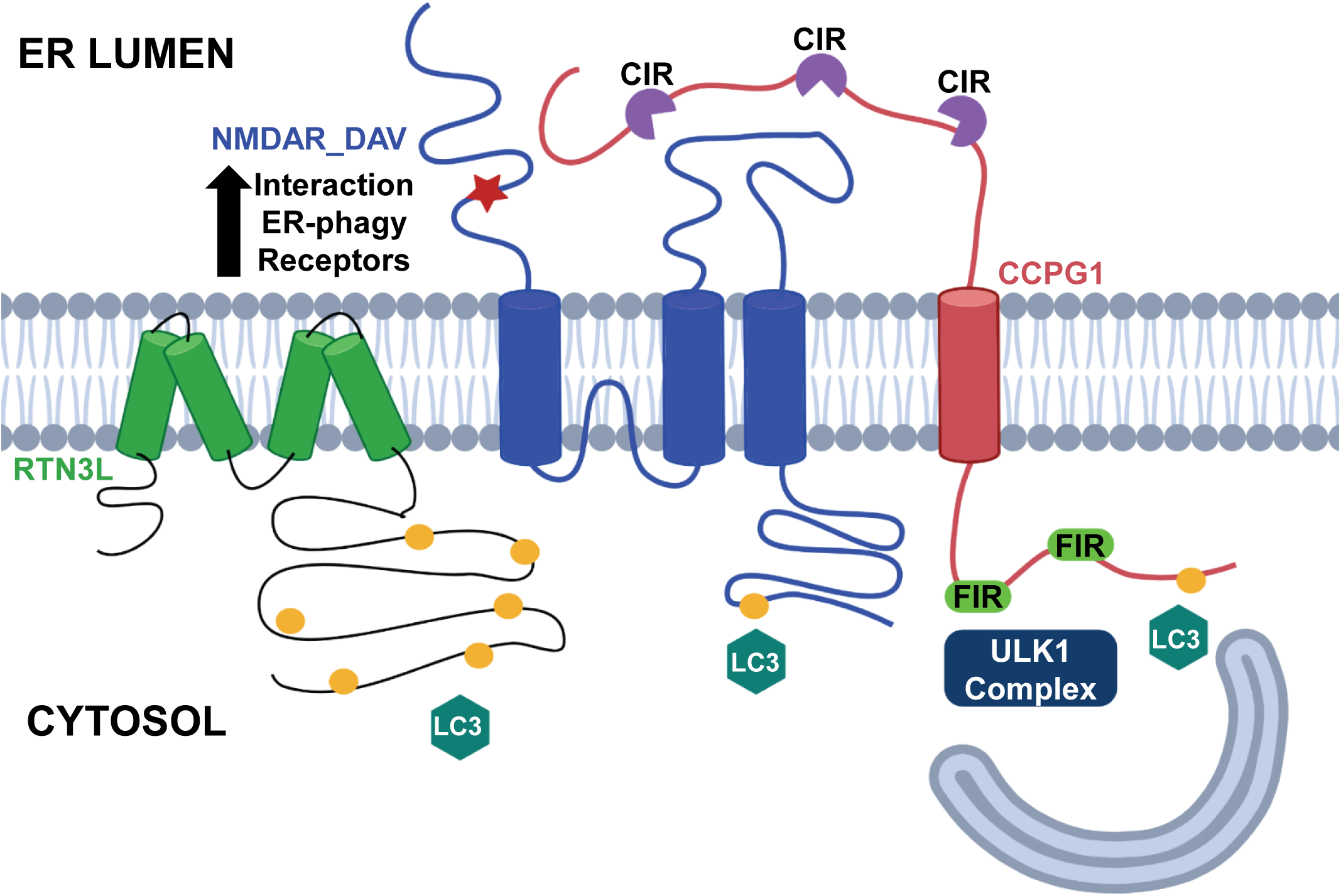
Proposed mechanism of autophagic clearance of the GluN2B R519Q variant. The GluN2B R519Q variant fails quality control measures in the ER resulting in its retention within the ER lumen. CCPG1, shown in red, present in the ER sheets is able to recognize the misfolded and accumulating R519Q variant, blue, and targets it for autophagic degradation via recruitment of autophagy machinery via interactions with the LIR, shown as yellow circles, and FIR domains, shown in neon green, of CCPG1. Additionally, RTN3L, shown in green, an ER-phagy receptor typically present in the ER tubules is able to interact with the R519Q variant, promoting its degradation through ER-phagy. It is also possible that the LIR domain present on the cytoplasmic CTD of the GluN2B subunit is able to interact with autophagy machinery directly.

An increasing body of work has investigated the effects of numerous missense mutations on the GluN2B subunit in various expression systems (Li et al., 2019; Platzer et al., 2017; Swanger et al., 2016; Vyklicky et al., 2018). A growing number of studies have implicated ligand binding to the GluN1, GluN2A, and GluN2B subunits as a quality control measure in the ER, perhaps serving to stabilize receptor subunits during assembly (Horak et al., 2014; Jeyifous et al., 2009; Lichnerova et al., 2015). Of particular interest, a recent study performed an alanine substitution at the R519 residue of the GluN2B subunit in order to investigate the early trafficking of NMDARs (Netolicky et al., 2024). They found that the R519A residue demonstrated a substantially reduced sensitivity to glutamate, with an EC_50_ of 1562 μM compared to the WT glutamate EC_50_ of 1.46 μM. This study also identified a strong correlation between the EC_50_ values of glycine binding with the GluN1 and surface expression of GluN1/GluN2A surface expression. Previous studies have also implicated motifs within the GluN1, GluN2A and GluN2B subunits that serve as ER retention signals that prevent trafficking of subunit prior to achieving their native conformation and assembly with other GluN subunits (Horak et al., 2008; Horak and Wenthold, 2009; Qiu et al., 2009; Scott et al., 2001; Standley et al., 2000; Xia et al., 2001). DAVs identified within the GluN2B subunit are particularly concentrated in the LBD (Swanger et al., 2016). Furthermore, NMDAR variants that have been characterized to have reduced surface expression are almost exclusively located in the LBD (Benske et al., 2022). For example, the E413G residue of the GluN2B within the LBD has been shown to promote the binding and unbinding of the glutamate ligand, resulting in reduced glutamate potency, despite not forming direct interactions with the glutamate it is an essential residue in the ligand binding pocket (Wells et al., 2018). Thereby, we focused on the highly uncharacterized variant, R519Q, and found that this substitution results in significantly reduced total and surface expression and stability of the GluN2B subunit (**Figure 1A-E**). Together, these results coincide with the report that the nonsense mutation, R519*, which results in the truncation of the GluN2B at this arginine residue, displays significantly reduced surface expression, a 50% decrease in peak current, and reduced spine density (Santos-Gómez et al., 2021).

The ubiquitin-proteasome and the autophagy-lysosome pathway are implicated in the clearance and degradation of misfolded, accumulated, and aggregated proteins in order to maintain the homeostasis of the cells. However, their precise involvement in the regulation of NMDARs has not been conclusively determined up until this point. Notably, upon treatment with Baf-A1 and MG132 to inhibit the lysosome and proteasome respectively, our data demonstrated that the R519Q was preferentially degraded by autophagy, as seen by the 4.2-fold accumulation upon Baf-A1 treatment (**Figure 2A**). However, it should be noted, under physiological conditions, it is likely the autophagy-lysosomal and ubiquitin-proteasome system work in tandem to regulate the proteostasis of NMDARs, given the equivalent accumulation upon inhibition of lysosomal and proteasomal degradation of the WT subunit (**Figure 2A, 5A**). The degradation of the R519Q variant via autophagy was recapitulated by pharmacological modulation of autophagy, in which autophagy inhibition resulted in the accumulation of the variant subunit (**Figure 2C**), and by depletion of autophagy genes by siRNA knockdown, in which knockdown of ATG7 and LC3b resulted in the accumulation of the R519Q variant (**Figure 2D, 2E**). Furthermore, co-immunoprecipitation demonstrated an interaction between LC3b and the WT and R519Q GluN2B subunit (**Figure 5E**). Implicating autophagy in the proteostasis of the GluN2B subunit was further validated by the overexpression of WT and R519Q variant GluN2B subunits in the autophagy deficient ATG7 KO HEK293T cells. Remarkably, the chronic loss of autophagy resulted in an increase in the R519Q variant, such that the total expression was equivalent to that of the WT (**Figure 3B**). Despite this, while depletion of autophagy increased the available subunits for assembly, it did not influence the surface expression of the R519Q variant and resulted in further accumulation within the ER (**Figure 3C**).

Further, we investigated the role of selective autophagy in the clearance of the R519Q variant in order to understand the specificity and efficiency. We discovered a highly conserved LIR motif within the GluN2B CTD (**Figure 4C**), which is not present in other GluN subunits (**Figure 4B**). Intriguingly, upon alanine substitution of the critical phenylalanine of this consensus sequence (**Figure 4A**), the LIR disruption restored the total expression of the R519Q_F1307A variant to that of the WT (**Figure 4D**). These results were similar to the total expression of the R519Q variant upon autophagy depletion in the ATG7 line (**Figure 3B**), indicating the LIR motif plays a critical role in the degradation of the R519Q variant. Nevertheless, disruption of the LIR domain was not sufficient to rescue the surface expression of the R519Q mutant in HEK293T or ATG7 KO cells (**Figure 4E, 4G**). Accordingly, disruption of the R519_F1307A LIR motif resulted in the greater accumulation of the GluN2B subunit after inhibition of the proteasome, shifting the efficient removal and degradation of the R519Q variant away from autophagy degradation (**Figure 5B**). Despite the accumulation of intracellular GluN2B, the inability of R519Q to traffic to the cell surface highlights a persistent defect in its folding or assembly that the increased expression as a result of autophagy inhibition cannot rescue. These results underscore the critical role of autophagy in the degradation of misfolded GluN2B subunits and reveal that R519Q degradation remains reliant on this pathway. Furthermore, they suggest that impaired autophagic flux does not suffice to overcome the intrinsic quality control barriers preventing mutant NMDAR surface expression. In addition, it is unlikely that chaperone-mediated autophagy plays a significant role, as the HSC70 adaptor protein is cytosolic, and consensus KFERQ-like motifs present in the GluN2B are in the ER lumen (Kirchner et al., 2019). However, it should not be entirely ruled out that posttranslational modifications such as acetylation and phosphorylation of the CTD of GluN2B subunits could target them to chaperone-mediated degradation.

It is known that single missense mutations can result in reduced efficiency of folding within the crowded ER lumen, leading to ER dysfunction which can activate the UPR. This has been demonstrated for membrane receptors such as GABA_A_Rs with frameshift variants in the α1 subunit(Williams et al., 2024). Future studies are needed to assess whether the R519Q variant leads to ER stress and ultimately activates the UPR, given its retention within ER compartments (**Figure 1F**). ER stress and the accumulation of damaged, misfolded, and aggregated proteins have been shown to induce their ubiquitination and degradation. It is thought that ER-associated degradation (ERAD) is responsible for the retrotranslocation and degradation of short-lived proteins through the ubiquitin-proteasome system. Certainly, it has been shown that inhibition of glycosylation in cortical neurons led to the degradation of GluN1 subunits (Gascón et al., 2007). Additionally, various E3 ligases, including NEDD4 (Gautam et al., 2013), Mib2 (Jurd et al., 2008), FBXO2 (Kato et al., 2005), and KTD13 (Gu et al., 2023), have been implicated in the ubiquitination of NMDAR subunits. However, to our knowledge, the direct degradation of NMDARs has not been thoroughly investigated in the context of the early secretory pathway. In this case, future studies could evaluate whether E3 ligases within the ER are responsible for maintaining the proteostasis of NMDARs. Until now, the degradation of the GluN2B receptor through autophagy has not been described. Conversely, it has been well demonstrated that the NMDAR can induce autophagy in neurons, specifically during long-term depression (LTD), in which NMDAR-mediated calcium signaling can influence the mTOR pathway, promoting autophagy, and driving the internalization and degradation of AMPA receptors through autophagy (Kallergi et al., 2022). However, it is presumed that, similarly to AMPARs, the endocytosis of NMDARs can result in lysosomal degradation via fusion with the endosome. Due to the accumulation within the ER and disposition towards autophagic degradation, we assessed whether ER-phagy was contributing to the turnover of R519Q variant GluN2B subunits. Knockdown of CCPG1 resulted in a striking increase in the GluN2B subunit. Additionally, the R519Q variant demonstrated enhanced interaction with CCPG1 compared to both WT and R519_F1307A LIR disrupted GluN2B subunits, and displayed a dose-dependent decrease upon overexpression with CCPG1 (**Figure 2F,2G**, **Figure 5E**). Furthermore, siRNA mediated knockdown of RTN3L (**Figure 2H**) resulted in the accumulation of the GluN2B. How these receptors engage the R519Q cargo merits further investigation.

Our previous work demonstrated a proof-of-concept model for remodeling the ER proteostasis in order to restore the surface expression and functionality of NMDARs. Specifically, we found that treatment with BiX, a potent BiP activator, modestly activates the IRE1 pathway of the UPR and enhances GluN2A variant folding, assembly, trafficking, and functionality (Zhang et al., 2024). However, studies by which the ER proteostasis network can be used to target and enhance the folding and assembly of variant NMDARs are severely limited by the lack of proteomic studies that provide insight into implicated proteins. While it is likely that well-studied chaperone proteins such as BiP and calnexin play a role in the biogenesis of NMDARs, the regional, functional, and temporal expression of NMDAR subunits indicates that there may be more specific pathways that regulate the expression of NMDARs. A large body of work investigating modulation of the ER proteostasis has focused on the inhibitory neurotransmitter receptor GABA_A_Rs. Recent work has identified HSP47, a heat shock protein with the capacity to correct the folding and incorporate variant α1 subunits into functional GABA_A_Rs via direct interaction and stabilization of the subunits (Wang et al., 2024). Additionally, work has demonstrated that these mutations can be rescued with a variety of proteostasis regulators and pharmacological chaperones, which bind to the protein and stabilize them during folding (Wang et al., 2022; Wang et al., 2023; Xi Chen, 2023). These methods to restore surface trafficking and functional signaling of receptors are novel and promising approaches to rescue trafficking-deficient mutations that display moderate loss-of-function phenotypes. Together, these findings highlight the significance of the work shown, beginning to identify potential targets for future therapeutics by understanding how these receptors are degraded under physiological and pathological conditions.

## Conclusion

In summary, this study demonstrates that the R519Q variant of the GluN2B subunit results in a loss-of-function of NMDARs and severely impacts their stability, expression, trafficking, and proteostasis. Furthermore, we identified that the R519Q variant drives the degradation of the GluN2B subunit towards autophagic degradation and clearance. Further characterization of the degradation mechanisms implicates the ER-phagy receptor CCPG1, as well as RTN3L, as playing a role in the removal of variant accumulations from the ER. Further, this study implicated the importance of the LIR motif that is conserved in the CTD in influencing the effectiveness of GluN2B autophagic degradation. Moreover, this study highlights that pathogenic DAVs within NMDAR subunits may exert their pathogenic phenotypes during early biogenesis, resulting in a loss of receptor function on the cell surface. We additionally revealed that the R519Q variant is subject to rigorous ER quality control measures that cannot simply be overcome to promote productive folding by preventing degradation and increasing the pool of variant subunits available for assembly. These findings may have broader impacts on other DAVs present within the LBD and nonsense mutations that result in receptor truncation. These findings provide the first look at the degradation and clearance of NMDARs through the autophagy pathway. This contributes new avenues of investigation into the functional rescue of NMDAR DAVs through targeted degradation of variant subunits, or through folding-correcting strategies by modulating adaptors of NMDAR proteostasis. This will allow researchers to identify new targeted therapies to regulate NMDAR function and expression.

### Limitations to the study

In this study, we identified the autophagy-lysosomal pathway is involved in the degradation of pathogenic GluN2B subunits on exogenously expressed NMDARs in HEK293T cells, using homozygous expression of variant subunits, while typically many patients present as heterozygous for these de novo variants. Future investigations are needed to validate the proteostasis of NMDARs in an endogenous environment and the role of the degradation pathways. Studies into the proteostasis of NMDARs are severely limited by the lack of knowledge in how these receptors fold and traffic within the ER membrane. Another remaining question is when and how the LC3b interacting motif plays a role in the degradation of this receptor subunit. Further investigations can further mutate the hydrophobic residues of this motif and see if there is an additional impact on the expression and degradation of receptor subunits.

## Materials and Methods

### Reagents

The cycloheximide (#ALX-380-269) was purchased from Enzo Life Sciences. The (+)-MK801 maleate (#HB0004) was obtained from HelloBio. MG132 (#A2585) and sulfo-NHS-SS-biotin (#A8005) were purchased from APExBio. EDTA-free protease inhibitor cocktail (#04693159001) was purchased from Roche. TransIT-2020 Transfection Reagent (#MIR5400) was obtained from Mirus Bio and the HiPerFect transfection reagent (#301707) was obtained from Qiagen Bovine serum albumin (BSA) (#A-421-25), G418 (G-418-10), and n-Dodecyl-β-maltoside (DDM) (#69227-93-6) were purchased from GoldBio. The 3-methyladenine (3-MA) (#HY-19312), SMER28 (#HY-100200), Rapamycin (#HY-10219), and DAPI dihydrochloride (HY-D0814) were obtained from MedChemExpress. The Bafilomycin-A1 (#11038) was obtained from Cayman Chemicals. Poly-L-lysine (#150177) was purchased from MP Biomedicals. All other chemicals were purchased from Sigma unless otherwise noted.

### Antibodies

The following primary antibodies were purchased and used in this study. The rabbit monoclonal anti-GluN2B antibody (#ab183942, 1:3000), rabbit polyclonal anti-GluN2B (#ab73001, 1:300), and rabbit monoclonal anti-Na^+^/K^+^ ATPase antibody (#ab76020, 1:20,000;1:300) were purchased from Abcam. The mouse monoclonal anti-GluN2B (clone N59/20, 75-097, 1:300) from NeuroMab was purchased from Antibodies Inc. The mouse monoclonal anti-β-actin (A1978, 1:10,000) was purchased from Sigma Aldrich. The rabbit polyclonal anti-p62 (PM045, 1:2000) was obtained from MBL. The rabbit monoclonal anti-CCPG1 (80158s, 1:2000), rabbit monoclonal anti-ATG7 (8558s, 1:1000), and rabbit monoclonal anti-Beclin 1 (3495s, 1:1000) came from Cell Signaling Technology. The rabbit polyclonal anti-LC3b (NB100-2220, 1:1500) was obtained from Novus Biologicals. The rabbit polyclonal anti-calnexin (ADI-SPA-860-F, 1:500) was purchased from Enzo Life Sciences. The mouse monoclonal anti-Ubiquitin (14-6078-82) was obtained from Invitrogen. The rabbit polyclonal anti-calnexin (10427-2-AP, 1:500) and rabbit polyclonal anti-RTN3 (12055-2-AP, 1:2000) were purchased from ProteinTech.

The following secondary antibodies were utilized for western blot detection: HRP conjugated goat anti-mouse IgG (H+L) Superclonal Recombinant antibody (Invitrogen #A28177, 1:10,000) and HRP conjugated goat anti-rabbit IgG (H+L) Superclonal Recombinant antibody (Invitrogen #A27036, 1:10,000). Alexa fluor secondary antibodies used for confocal immunofluorescence are as follows: Alexa fluor 488 goat anti-mouse antibody (Invitrogen #A11029), Alexa fluor 488 goat anti-rabbit antibody (Invitrogen #A11034), Alexa fluor 568 goat anti-mouse antibody (Invitrogen #A11031), and Alexa fluor 568 goat anti-rabbit antibody (Invitrogen #A11036).

### Plasmids and Mutagenesis

The pcDNA3.1-GRIN1 (OHu22255D, NM_007327, human) and the pcDNA3.1-GRIN2B (OHu26128D, NM_000834, human) and pcDNA3.1-CCPG1 (OHu07897C, NM_004748.5) were obtained from GenScript. The R519Q and F1307A mutations were introduced into the GRIN2B plasmid using QuikChange II site-directed mutagenesis kit (Agilent Genomics, #200523). All cDNA sequences were confirmed using DNA sequencing.

### Cell Culture and Transfection

HEK293T cells (ATCC, #CRL-3216 or Abgent, #CL1032) were maintained in Dulbecco’s Modified Eagle Medium (DMEM) (Cytiva #SH30243.01) containing 10% heat-inactivated fetal bovine serum (Cytiva, #SH30396.03HI) and 1% penicillin streptomycin (Cytiva #SV30010) at 37°C in 5% CO_2_. Maintenance plates were passaged using 0.05% trypsin protease (Cytiva, #SH30236.01) upon reaching no greater than 90% confluency, and were used for experiments from passages 5-30. Cells were seeded in 10-cm dishes, 35-mm dishes or 6-well plates and grown until reaching 50-70% confluency. Confluent cells were transiently transfected with a 1:1 ratio of GluN1:GluN2B plasmids using TransIT-2020 Transfection Reagent (Mirus Bio #MIR5400) according to the manufacturer’s instructions. In order to prevent glutamate-mediated excitotoxicity, the media was supplemented with 50 µM (+)-MK-801 (HelloBio #HB0004) and 2.5 mM MgCl_2_ (Sigma, #208337) 4 hr post-transfection. 48 hr post-transfection, cells were harvested for protein analysis.

Stable HEK293T (Abgent, #CL1032) cell lines expressing NMDARs composed of GluN1_GluN2B and GluN1_GluN2B(R519Q) were generated using a G418 (Enzo Life Sciences, #ALX-380-013-G001) selection method in the presence of channel blockers MK-801 and MgCl_2_ as previously described. Briefly, cells were transfected with a 1:1 cDNA ratio of GluN1:GluN2B and GluN1:GluN2B(R519Q) respectively. Cells were selected using DMEM supplemented with 0.8 mg/mL G418 for 14 days. Cells were then maintained in DMEM supplemented with 0.4 mg/mL G418 and expression of GluN1 and GluN2B subunits were verified using western blot analysis.

### Discovering the LIR motif and Sequence Alignment

In order to demonstrate the involvement of the LIR motif, the complete amino acid sequence of the GRIN2B gene was obtained by assessing the UniProt Database (Q13224). A search was performed using the iLIR web server (https://ilir.warwick.ac.uk/) to identify peptide sequences associated with the LIR motif. Within the amino acid sequence of GRIN2B, a canonical LIR fragment, DTFVDL, was identified at positions 1305-1310. Sequences of the Homo Sapiens (human), Mus musculus (mice), Rat norvegicus (rat), Canis lupus familiaris (dog), and Pan troglodytes (chimpanzee) GluN2B subunits were obtain through Uniprot Database (respective accession Numbers: Q13224, Q01097, Q00960, Q5R1P3, H2Q5I0). Sequence alignment was performed using Clustal Omega.

### RNA interference (RNAi)-mediated gene knockdown

HEK293T cells stably expressing R519Q_GluN2B NMDARs were grown until they reached 60% confluency. 50 nM of siRNA was transfected using HiPerfect Transfection Reagent (Qiagen # 301717) according to the manufacturer’s instructions. 48 hr after transfection, cells were harvested and subjected to SDS-PAGE and western blot analysis. Two distinct siRNAs were used for each gene knockdown, independently and in combination, in order to minimize off-target effects and increase knockdown efficiency. Non-targeting scrambled siRNA was used as a control (Dharmacon #D-001810-01-20).

The following human ON-TARGET*plus* siRNA duplexes were obtained from Dharmacon: CCPG1.1 (J-013998-05-0005), CPPG1.2 (J-013998-06-0005), CCPG1.3 (J-013998-07-0005), CCPG1.4 (J-013998-08-0005), RTN3.1 (J-020088-09), RTN3.2 (J-020088-10), ATG7.1 (J-020112-05-0005), ATG7.2 (J-020112-06-0005), LC3a.1 (J-013579-07-0005), LC3a.2 (J-013579-08-0005), LC3b.1 (J-012846-05-0005), LC3b.2 (J-012846-08-0005). LC3c.1 (J-032399-09-0005), and LC3c.2 (J-032399-11-0005). Non-targeting (NT) control pool siRNA (#D-001810-01-20) was used as a negative control. The designation of GENE.1 and GENE.2 indicates two distinct siRNA sequences against each mRNA transcript.

### SDS-PAGE and Western Blot

Cells were harvested using 4°C Dulbecco’s phosphate-buffered saline (DPBS) (Corning #21-030-CVR) and lysed using lysis buffer (50 mM Tris-HCl, pH 7.4, 150 mM NaCl, 2 mM n-Dodecyl-β-D-maltoside (DDM) (GoldBio, #DDM5)) supplemented with protease inhibitor cocktail (Roche, #04693159001). Lysates were cleared via centrifugation (20,000 xg, 10 min, at 4°C) and the supernatant was collected as total protein. Protein concentrations were measured using the MicroBCA Protein Assay Kit (ThermoFisher, #23235). 30 µg of cell lysates were loaded with Laemmli buffer (Biorad #1610747) with 10% β-mercaptoethanol (βME) (Sigma #M6250). Samples were subjected to SDS-PAGE and separated on either an 8% or a 4-20% acrylamide gel. The Precision Plus Protein Kaleidoscope Pretained Protein Standard (Biorad #31610375) and/or the Broad Range Prestained Protein Marker (Proteintech #PL00002) were used as standards. Western blot analysis was performed, and gels were transferred to 0.45 µm nitrocellulose membrane (Biorad #1620115) and then blocked in 5% skim milk in TBST buffer for 1 hr at room temperature. Membranes were probed with the appropriate antibodies and dilutions listed above. β-actin and Na^+^/K^+^ ATPase were used as loading controls for total protein lysate and plasma membrane proteins respectively. Membrane visualization was performed using chemiluminescent substrates SuperSignal West Pico PLUS Chemiluminescent Substrate (ThermoFisher #34578) or SuperSignal West Femto Maximum Sensitivity Chemiluminescent Substrate (ThermoFisher #34096) and images were acquired using Azure 600 Imager (Azure Biosystems). Protein band intensity was quantified using Image J software from the NIH. Total protein was normalized to the loading control and then the experimental control (DMSO vehicle or WT).

### Biotinylation of Cell Surface Proteins

HEK293T cells were plated on poly-L-lysine coated plates and cultured as described. Surface biotinylation experiments were performed according to published procedures (Wang et al., 2022; Zhang et al., 2024). Briefly, cells were transfected with the corresponding GluN1 and GluN2B plasmids for 48 hr. If being treated with an autophagy regulator, cells were treated 24 hr after transient transfection for 24 hr. Intact cells were washed with ice-cold DPBS and incubated with 0.5 mg/mL sulfo-NHS SS Biotin (APExBio #A8005), a membrane impermeable biotinylation reagent, dissolved in DPBS containing 1 mM CaCl_2_ and 0.5 mM MgCl_2_ (DPBS+CM) for 30 min at 4°C to label surface membrane proteins. Cells were then twice incubated with 50 mM glycine in DPBS+CM for 5 min at 4°C to quench the labeling reaction. Sulfhydryl groups were blocked by incubating the cells with 5 nM N-ethylmaleimide (NEM) for 15 min at room temperature. Cells were scraped off in lysis buffer (50 mM Tris-HCl, pH 7.4, 150 mM NaCl, 2 mM DDM) supplemented with 5 mM NEM and protease inhibitor cocktail. Cells were solubilized overnight at 4°C and lysates were cleared by centrifugation at (20,000 xg, 10 min at 4°C). The concentration of the supernatant containing the biotinylated surface proteins was measured using a MicroBCA assay. Biotinylated surface proteins were affinity-purified by incubating the supernatant with 70 µL NeutrAvidin-conjugated agarose beads (ThermoFisher #29201) at 4°C for 18-24 hr. The samples were subjected to centrifugation (5,000 xg, 1 min, at 4°C). The beads were washed 6X with solubilization buffer (1X TBS, 1% Triton X-100). Surface proteins were eluted from beads by vortexing for 20 minutes with 120 µL of elution buffer (2X Laemmli sample buffer, 100 mM DTT, 6M urea pH 6.8) followed by SDS-PAGE and western blot analysis.

### Cycloheximide-Chase Assay

HEK293T cells stably expressing GluN1_GluN2B, and GluN1_GluN2B(R519Q) NMDAR receptors were treated with 100 µg/mL cycloheximide (Enzo Life Sciences, #ALX-380-269-G001) to inhibit protein translation and synthesis. Cells were chased for the indicated amount of time, harvested, and lysed for SDS-PAGE and western blot analysis to assess total protein.

### Confocal Immunofluorescence Staining and Confocal Microscopy

HEK293T cells were fixed with 4% paraformaldehyde in DPBS for 20 min and then blocked with 3% BSA in DPBS for 30 min. For surface staining, cells were incubated with 3% BSA containing mouse-monoclonal anti-GluN2B (1:300) and rabbit monoclonal anti-Na^+^/K^+^ ATPase antibody (1:300) overnight at 4°C, without detergent permeabilization. To investigate accumulation in the ER, membranes were permeabilized by incubating the fixed cells with 0.2% saponin for 20 min at room temperature prior to blocking. The cells were then incubated with 4% BSA containing mouse-monoclonal anti-GluN2B (75-097, 1:300), rabbit polyclonal anti-Calnexin (1:500), and 0.2% saponin overnight at 4°C. The cells were washed thoroughly with DPBS after primary staining and incubated with Alexa 488-conjugated goat anti-mouse antibody (1:500 and Alexa 568-conjugated goat anti-rabbit antibody (1:500) diluted in 3% BSA for 1 hr at room temperature. Cell were permeabilized with 0.2% saponin and incubated with 1 µg/mL DAPI for 10 min. The coverslips were then mounted with Fluoromount-G (Invitrogen, #00-4958-02) and sealed. An Olympus IX-81 Fluoview FV3000 confocal laser scanning microscope was used. A 60X oil objective was used to collect high-resolution images using FV31s-SW software. The images were analyzed using Image J software.

### Immunoprecipitation

Cell lysates (1000 µg) were pre-cleared with 30 µL of Protein A/G plus-agarose beads (Santa Cruz Biotechnology, sc-2003) and 1 µg normal mouse IgG (Santa Cruz Biotenchnology, #sc-2025) for 1 hr at 4°C to remove nonspecific binding proteins. The pre-cleared lysates were incubated with 2 µg of mouse anti-GluN2B antibody for 1 hr at 4°C, and then 50 µL of Protein A/G-plus agarose beads were added and incubated overnight at 4°C. The beads were collected by centrifugation at 8000 xg for 1 min and washed 3X with lysis buffer. Protein complexes were eluted by incubating with 100 µL of Laemmli sample buffer with BME. The eluents were subjected to SDS-PAGE on a 4-20% gradient gel and western blot analysis was performed. IgG serves as a negative control.

### Quantification and Statistical Analyses

All data was analyzed using GraphPad Prism software. All data are presented as mean ± SEM. Statistical significance was evaluated between two groups using two-tailed Student’s t-test or a Mann Whitney test for nonparametric data. Comparison between multiple groups, a one-way ANOVA followed by a post-hoc Tukey or Dunnett’s post hoc test for comparisons. An assigned p-value < 0.05 was considered statistically significant. *, p<0.05; **, p<0.01; ***, p<0.001; ****, p<0.0001.

## Data availability

All data are contained within the manuscript.

## Online supplemental material

This manuscript contains one supplemental figure.

## Conflict of interest

The authors declare that they have no conflicts of interest with the contents of this article.

## Acknowledgements

We would like to thank Dr. Boaz Tirosh (Case Western Reserve University, Cleveland, Ohio) for the HEK293T ATG7 KO cell line used in these studies. This work was supported by the National Institutes of Health grant R01NS117176 awarded to T. Mu.

## Author contributions

T. Benske: Conceptualization, Data curation, Investigation, Methodology, Formal Analysis, Visualization, Validation, Writing – original draft, Writing – review and editing, M. Williams: Data curation, Investigation, Methodology, Writing – review and editing, P. Zhang: Writing-review and editing, A. Palumbo Visualization, Software, Writing – review and editing, T. Mu Conceptualization, Formal Analysis, Visualization, Resources, Funding acquisition, Project administration, Supervision, Validation, Writing – review and editing.

---

**Supplementary Figure 1:**
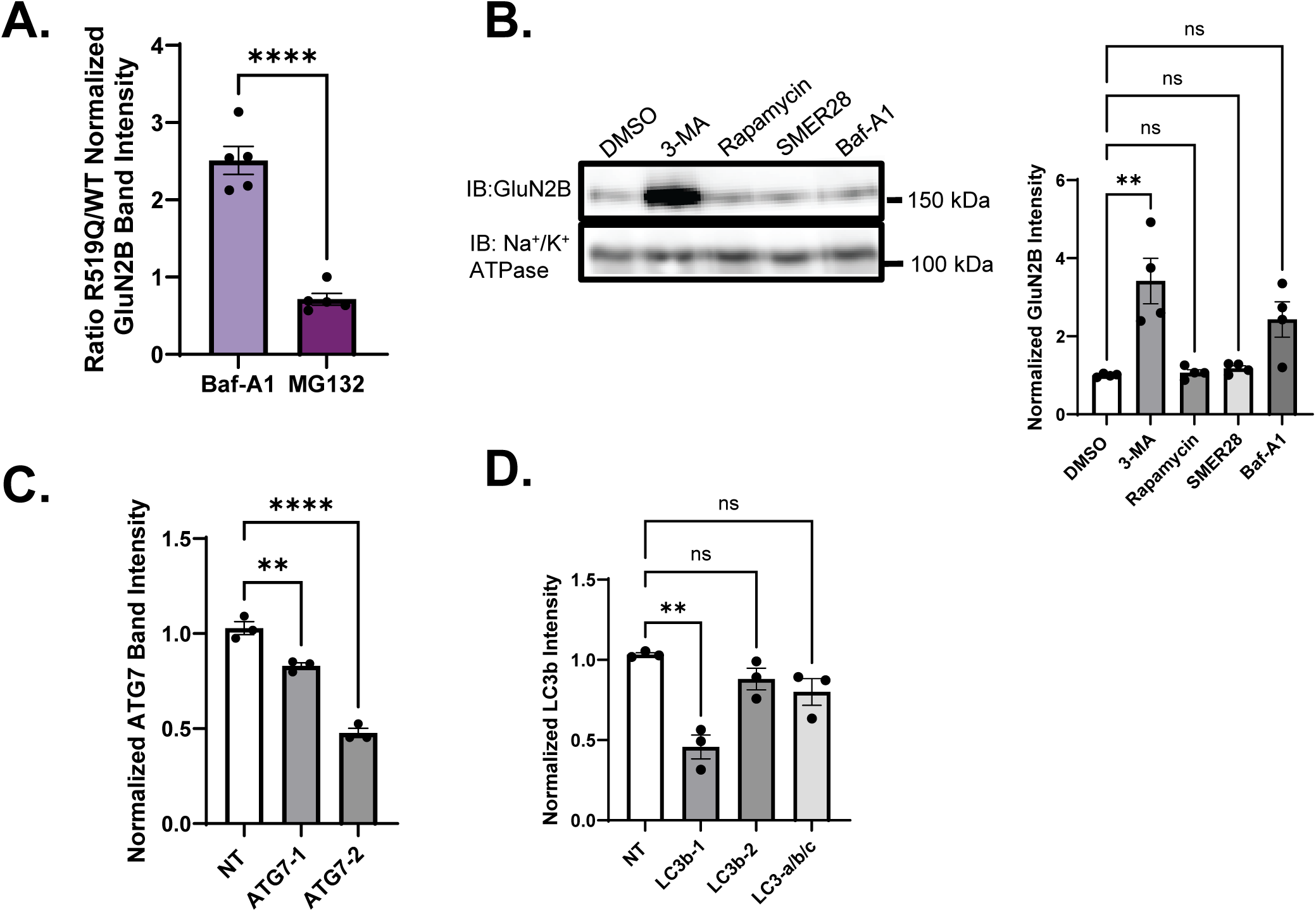
Effects of autophagy modulation on the R519Q variant. **(A)** Ratio of the normalized accumulation of R519Q/WT GluN2B upon treatment with Baf-A1 and MG132 from Figure 2A. **(B)** HEK293T cells stably expressing R519Q NMDARs were treated with autophagy activators (SMER28 10 µM, Rapamycin 100 nM) and inhibitors of autophagy (3-MA 50 mM and Baf-A1 20 nM) for 24 hours. Surface biotinylation assay to monitor the influence on the surface expression 24 hrs after drug treatment (n=3). Na^+^/K^+^ ATPase served as a membrane protein loading control. **(C)** Quantification of the ATG7 siRNA knockdown efficiency (n=3). β-actin served as the soluble total protein loading control. **(D)** Quantification of the LC3b siRNA knockdown efficiency (n=3). β-actin served as the soluble total protein loading control.

## Notes

### Competing Interest Statement

The authors have declared no competing interest.

